# Simulating Tandem Mass Spectra for Small Molecules using a General-Purpose Large-Language Model

**DOI:** 10.1101/2025.11.10.687298

**Authors:** Tuan Nguyen, Dinesh Barupal

**Author notes:** Corresponding author: Address: CAM Building, 3rd floor, 17 E 102nd St, New York, NY 10029.

## Abstract

We show a practical application of the Google Gemini large-language-model for simulating tandem mass spectra for compounds from the Blood Exposome Database. This approach bypasses the need for domain-specific model training, suggesting that the chemical fragmentation knowledge could be latently encoded within the Gemini model. General-purpose LLMs represent a useful and accessible tool for expanding *in-silico* spectral libraries and may accelerate the compound annotation in mass spectrometry-based metabolomics and exposomics.

## Introduction

A majority of compounds in the metabolomic and exposomic knowledgebases lack mass spectral data^1^, limiting their use in annotating high-resolution mass spectrometry datasets. Collecting experimental data for these compounds (n > 100,000) is cost-prohibitive, but in-silico approaches^2^ can predict mass spectra for them. For that, three types of approaches exist, rule-based^3^, quantum chemistry^1,4^ and machine learning^2,5^. Additionally, the new deep learning architecture, transformer underling the large-language models (LLMs) which have been driving the recent AI revolution, can be used for predicting in-silico spectra^6^. Large-language models have shown practical applicatiions in biomedical research to summarize clinical notes^7,8^, predict protein structures^9^, estimate binding affinities^10^, and predict biochemical reactions^11,12^. Here, we report that Google’s Gemini model, which is generally available for anyone to use via their Application Programming Interfaces (APIs) and online interface (gemini.google.com), can predict highly accurate mass spectra for small molecules, effectively bypassing the need to train a new LLM that translate structures to mass spectra. We hypothesize that these LLMs, with their vast pre-trained knowledge of chemical principles and their ability to perform complex reasoning, can predict mass spectra in a zero-shot or few-shot setting.

## Results and discussion

We optimized a chain of thought (CoT) LLM prompt (<u>Zenodo Accession</u>) for SMILE code, monoisotopic mass, adduct type, and chemical name as input and asked the return Gemini model t the mass spectra (M+H) in the NIST MSP format.

To our surprise, Gemini returned a highly accurate mass spectra (Figure 1) for the initially tested compound, Acetaminophen, a commonly used pain-killer medicine (Figure 1). It correctly returned all the major m/z peaks in the mass spectra and their intensities that are comparable to the true positive spectra in the NIST 2023 database (Figure 1). Excited by this tryout, we further tested the Gemini LLM for predicting M+H mass spectra for 1436 compounds from the Blood Exposome Database^14^, that have mass spectra available in the NIST23 database. We observed that 62% of tested compounds showed a higher similarity (Cosine Score > 0.70, with 3 or more matching peaks) between their Gemini simulated mass spectra and the NIST 23 spectra (Figure 2) (<u>Zenodo Accession</u>). We also predicted negative mode (M-H) mass spectra for 835 compounds (<u>Zenodo Accession</u>) from the Blood Exposome Database. Of the tested compounds, 0.53 % had a Cosine score of 0.70, with 3 or more matching peaks (Figure 2). For many other low scored spectra, several neutral losses were correctly predicted by the Gemini LLM. It took less than one hour to generate these mass spectra for both modes using the parallel use of the Gemini live API. We did not notice a bias towards any chemical class or sub-structure (Figure 2), also the repeated testing (n=20) for the same compound yielded almost identical spectra. These results suggested that Gemini model can be used for creating large-scale in-silico mass spectral libraries.

**Figure 1.**
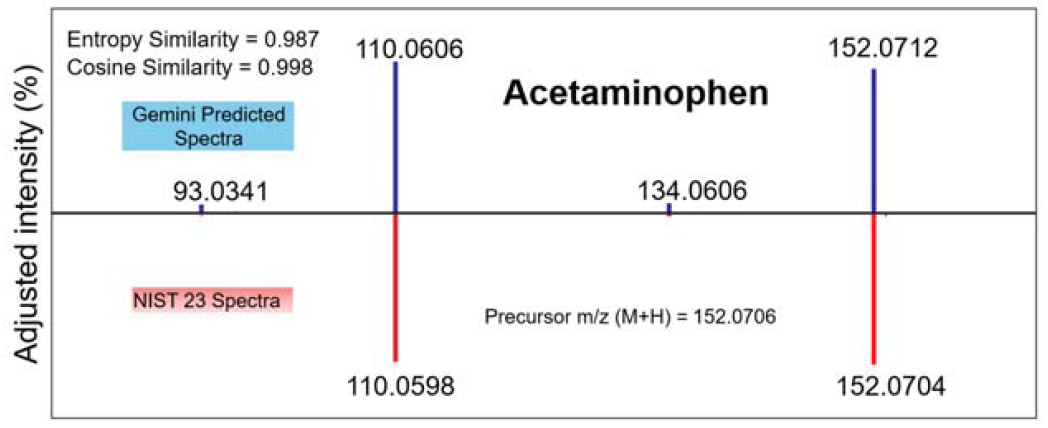
A general-purpose large language model predicted high-quality mass spectra for a small molecule. We used Google’s Gemini LLM model for the prediction of mass spectra. IDSL.CSA software^13^ was used for computing the similarity scores and the draw the spectra similarity plot.

**Figure 2.**
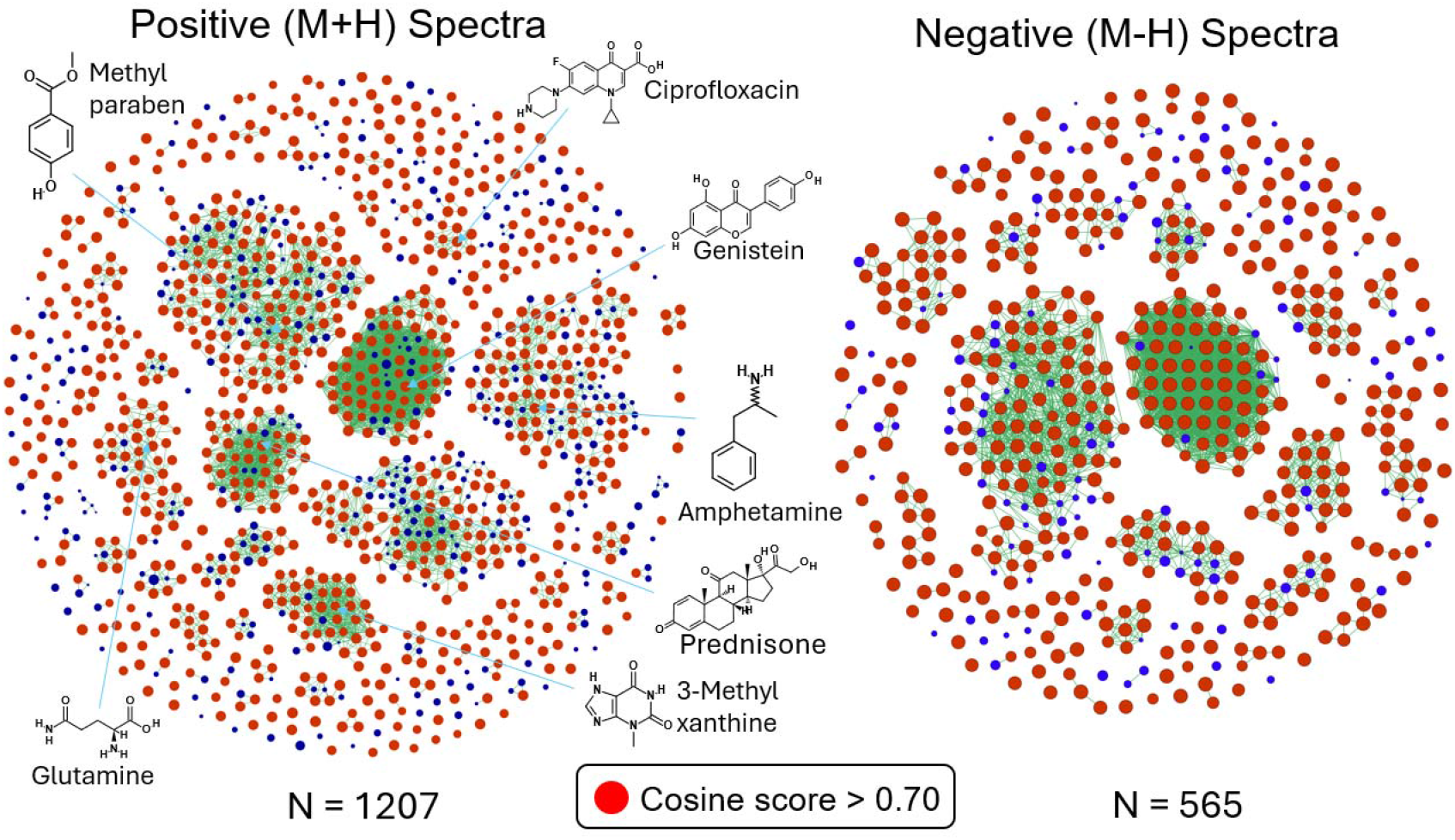
Gemini can predict spectra for a wide diversity of chemicals. These chemical similarity networks highlight the results of the spectral similarity between the Gemini predicted spectra and the NIST 23 spectra for a compound (node). The edge between nodes shows a Tanimoto score of 0.75 and node size is proportional to the cosine score; largest node has 1.0 score. Compounds for which in-silico and NIST23 spectra had at least 3 overlapping m/z peaks were included.

The accurate prediction of the mass spectra can be attributed to the mixture of expert and chain-of-thought advancements in the Gemini LLM. The Mixture-of-expert(MoE) architecture^15,16^ which allows LLM to accumulate specialized similar/related domain specific topics knowledge during training by assigning data of similar topics to a subnetwork of the model, called an expert, that develops non-overlapping and focused knowledge based on the assigned topics. It is likely that Gemini training data contains significant amount of material related to mass spectrometry domain, which the MoE architecture and CoT prompt method can take advantage of during spectra prediction.

We advocate a practical use of these generally available online LLM models for expanding mass spectral libraries for small molecules.

## Methods and materials

Pre-trained LLM models can be run locally or accessed by an API offered by commercial providers such as Google or OpenAI. Implementing a local model is quite resource demanding due to the large number of parameters, so utilizing the online version via an API is preferred for many predictive tasks such as curating a chemical list or summarizing clinical importance of a gene list. Moreover, the Gemini model is available via the online site gemini.google.com and is estimated to be used by over 350 million users per month and is freely available to many academic institutions. This wide accessibility of the Gemini LLM model can democratize the mass spectra prediction and promote a major expansion of mass spectral libraries.

### Google Gemini Prompt Engineering

We engineered an optimized prompt to instruct Gemini LLM to generate the mass spectra for an input compound. The prompt requires SMILES, Chemical Name, Monoisotopic Mass and Molecular Formula as input, which can be obtained from the PubChem Database. The chain-of-thought prompt is provided at (<u>Zenodo Accession</u>). This long-form prompt was designed to guide the model’s reasoning process. This prompt instructed the model to find likely fragmentation mechanisms and stable neutral losses, and then to construct the spectrum based on that reasoning. The quality of the predicted spectra was highly dependent on the prompt design. The chain of thought (CoT) prompts^17^ consistently produced spectra that were more chemically plausible and achieved significantly higher similarity scores compared to the simple prompts. Examining model’s thinking summary confirms its ability to comprehend and follow detailed guidelines when predicting spectra. We also assessed reproducibility by running predictions for a subset of 100 molecules three times each. The determinism of the model was tested by setting the API’s temperature parameter to zero with fixed seed, which resulted in identical outputs for identical inputs, confirming high reproducibility under controlled settings. However, upon further testing, we noticed the quality of generated spectra under more deterministic condition is slightly lower compared to the default setting set by Gemini (default temperature = 1.0). A potential explanation is at temp=0, the model always selects the most probable fragments, which might create overly predictable spectra that miss important but less common fragmentation pathways. Real MS/MS spectra often contain minor peaks from alternative fragmentation routes that are chemically valid but statistically less frequent in training data. The main trade-off for using higher temperature parameter is lower reproducibility, as the same input can results in different but highly similar output variants. The Gemini agent was provided with a system-level instruction set prior to receiving any prompts. This instruction defined its role, output format, and constraints. For example, it was instructed to always return the spectrum as a JSON object containing two lists (m/z values and corresponding relative intensities) to ensure consistent, machine-readable output.

### Blood Exposome Database

To test our method on a chemically diverse and biologically relevant set of molecules, we utilized the Blood Exposome Database^14^ (www.bloodexposome.org), also available at https://zenodo.org/records/8146024. This database contains thousands of compounds that have been reported for blood specimens, representing a wide range of chemical classes, including metabolites, environmental contaminants, and food components. The canonical SMILES string for each compound was extracted to serve as the input for our prediction workflow. The 5000 top compound, as sorted by the literature count were selected. InChiKey for these compounds were against the NIST23 dataset for checking the availability of [M+H]+ and [M-H]− adducts mass spectra. A total 1436 InChiKey with matching spectra in [M+H]+ and 835 InChiKey with spectra in [M-H]−. Chemical properties, including SMILES, Monoisotopic mass and Molecular Formula were collected using PubChem’s PUGREST API (https://pubchem.ncbi.nlm.nih.gov/docs/pug-rest) for each InChiKey.

### Gemini Agent API

All predictions were generated programmatically using the Gemini API. A Python script was developed to iterate through the list of compounds from the Blood Exposome Database, format the appropriate prompt for each, and send requests to the API. The structured spectral data was then parsed from the model’s response and stored for analysis. The final results were exported to the NIST MSP format. These results are available at (<u>Zenodo Accession</u>).

#### Validation Against the NIST23 Library

The primary validation was performed by comparing the Gemini-predicted spectra against the gold-standard experimental spectra from the NIST23 library. For each compound, the cosine similarity and entropy similarity score were calculated between the predicted and experimental spectrum using the IDSL.CSA software^13^. The parameters files for the spectra similarity searches are made available at (<u>Zenodo Accession</u>). The results show that the chain of thought (CoT) prompt strategy achieved a mean cosine similarity score that is competitive with some dedicated ML models, demonstrating the viability of LLMs for this task even without specific fine-tuning.

## Author’s contribution

TN and DB planned the study, implemented the scripts, prepared the results and drafted the manuscript. All authors have reviewed the manuscript’s content.

## Data and code availability

Data and code are available at https://zenodo.org/records/17555571

